# Safeguarding *Iguana* diversity: Enabling rapid and low-effort tracking of non-native iguanas through terrestrial eDNA innovations

**DOI:** 10.1101/2024.11.28.625859

**Authors:** Jeroen L. van Kuijk, Matthijs P. van den Burg, Emilie A. Didaskalou, Mark de Boer, Adolphe O. Debrot, Ben Wielstra, Kathryn A. Stewart

**Author notes:** Corresponding Author: Dr. K. Stewart.

## Abstract

Reptiles have among the highest extinction risk across terrestrial vertebrates, with habitat fragmentation, habitat destruction, and invasive alien species being the primary causes of reptile species loss on a global scale. Invasive hybridization (i.e. hybridization between native and invasive alien species) is increasing globally, causing the extinction of native genotypes, and this phenomenon is particularly pervasive in Caribbean iguanas. The Lesser Antillean Iguana (*Iguana delicatissima*), a keystone species of Caribbean coastal ecosystems, has become critically endangered mainly due to ongoing hybridization with the invasive Common Green Iguana (*I. iguana*). For impactful conservation intervention, the need for early detection of invasive animals and their progeny, or detection of surviving pure native animals, is urgent. We aimed to develop a novel environmental DNA (eDNA) toolkit using Kompetitive Allele Specific PCR (KASP) technology, a method of allele-specific amplification for cost-effective and efficient sampling of terrestrial substrates to aid in mapping the distribution of native *I. delicatissima*, invasive *I. iguana*, and signal potential invasive hybridization. We demonstrate proof-of-concept and successfully identified *I. delicatissima, I. iguana,* and their hybrids via blood samples using our primer sets, as well as successful detection of *I. delicatissima* in several ex-situ (Rotterdam Zoo) and in-situ (St. Eustatius) eDNA samples, collected with environmental swabs and tape-lifting. We found that sampling potential perching spots yielded the highest number of positive detections via environmental swabbing and tape-lifting. Our toolkit demonstrates the potential of terrestrial eDNA sampling for iguana conservation, enabling faster detection of potential invasive hybridization. Additionally, the method holds promise for other terrestrial cryptic species, contributing to broader collection of population-level information.

## Introduction

Reptiles have among the highest extinction risk of all terrestrial vertebrates, with habitat fragmentation, destruction, and invasive alien species (IAS) being the primary causes of global species loss (Cushman 2006; Cordier et al. 2021). Of the approximate 70% decline in global vertebrate populations, indirect and direct translocation, introduction, and diffusion of IAS represent a major contributor to biodiversity decline (Bellard et al. 2016). Importantly, IAS not only endanger wildlife via predation, competition, and disease transmission (Ricciardi et al. 2017), they also threaten the genetic integrity of native taxa through hybridization (Meilink et al. 2015; van den Burg et al. 2023). In fact, invasive hybridization (hybridization between native and IAS) is underappreciated, but significant cause of species decline (Vuillaume et al. 2015), leading to the extinction of native genotypes, eliminating local adaptations or entire species and ultimately homogenizing biodiversity (Largiadèr 2008). The consequences of invasive hybridization can thus have far-reaching consequences, particularly when native individuals show lower fitness than their hybrid counterparts (Prentis et al. 2008). Therefore, for impactful conservation intervention, the need for early detection of hybridization or, depending on the stage of invasion, detection of (presumably rare) surviving pure native animals, is more urgent than ever.

One of the many indigenous and enigmatic insular species groups threatened by invasive hybridization are West Indian iguanas (Alberts et al. 2004). Herbivorous reptiles such as iguanas perform fundamental ecosystem services via herbivory, nutrient cycling, and seed dispersal of indigenous plants (Cooper and Vitt 2002; Traveset et al. 2016; Valido and Olesen 2019). The role of iguanas as keystone-species highlights the need for the protection of dry coastal and elevated coastal forests with their corresponding communities, as these threatened environments also harbor several other endemic species. Preservation of these communities for iguanas will in-turn contribute to the mitigation of the dramatic loss of biodiversity in the Caribbean (Knapp et al. 2014; Thibaudier et al. 2024).

The Caribbean Lesser Antilles are inhabited by high iguanid diversity, though these native populations can still hybridize with non-native iguanas (Stephen et al. 2013). In particular, the critically endangered Lesser Antillean Iguana (*Iguana delicatissima*) is at risk of hybridizing with non-native Common Green Iguanas (*Iguana iguana*), whilst its populations are also threatened by other anthropogenic factors (van den Burg et al. 2018a). *Iguana delicatissima* represents a unique component of Caribbean biodiversity, though only five populations remain that are free of the immediate threat of hybridization with non-native iguanas (van den Burg et al. 2018a; Pounder et al. 2020). St. Eustatius is home to one of the few remaining and very small *I. delicatissima* populations, with a recent estimate of ∼1000 individuals (van den Burg et al. 2022). The compounded effect of invasive green iguanas and their ensuant hybridization on St. Eustatius is complicating population recovery (van den Burg et al. 2018b).

Displacement by, and hybridization with, *I. iguana* are the dominant factors in the decline of *I. delicatissima* across its range, with few island populations remaining that are not directly threatened by invasive *I. iguana* (Pounder et al. 2020). These negative processes have already led to the extinction of several *I. delicatissima* populations, e.g., several islands within the Guadeloupe Archipelago and in southern Martinique (Vuillaume et al. 2015). In fact, not only is identification and removal of invasive iguanas directly needed, the speed and spread of invasive hybridization is a compounding practical conservation hurdle. Hybridization initially poses a less visible threat, but quickly escalates once the invasive population grows. Such rapid effects to *I. delicatissima* population sustainability and recovery prompts immediate choices for impactful detection and extraction.

Commonly used methods for individual detection and population monitoring are transect-based morphological surveys. These conventional surveys, however, often have low detection rates when applied to rare or cryptic species (Crawford et al. 2020; Matthias et al. 2021). Conventional morphologically-based surveys often require hand-capture and individual genotyping, causing risk to caught organisms, as well as much time and cost for accurate processing (Stewart and Taylor, 2020). Importantly, morphologically-based surveys are also particularly unreliable for identifying hybrid individuals. Breuil (2013) and Vuillaume et al. (2015) have shown that even animals identified in the field as *I. delicatissima* were sometimes hybrid or introgressed individuals, and although recent research (van den Burg et al. 2024) has identified additional morphological characteristics that might be used to detect hybrids, detailed data needed for its application and a study on its broader utility remain lacking. Since morphology still does not provide identification certainty, there remains a need for alternative monitoring techniques for the detection of rare invasive or hybrid individuals. This is particularly true early during invasion before they spread and multiply, when eradication efforts are likely to have some chance at success (e.g., Debrot et al. 2022). Similarly, the detection of rare surviving native animals for removal to artificial breeding programs has high conservation value (e.g., Milinkovitch et al. 2013; Miller et al. 2017; Pounder et al. 2020).

The application of environmental DNA (eDNA) is a well-suited and powerful, yet relatively underused tool in terrestrial biodiversity monitoring (Banerjee et al. 2022; Lyman et al. 2022; Wilcox et al. 2022; Aucone et al. 2023; Lynggaard et al. 2024). This is surprising considering it is non-invasive, non-destructive, time and cost efficient, and eliminates the considerable sampling effort and animal-handling expertise often needed with conventional methods (Stewart et al. 2017; Qu & Stewart. 2019; Didaskalou et al. 2022). Fortunately, sampling of the terrestrial environment for reptile eDNA is a growing field of research (Hunter et al. 2015; Kucherenko et al. 2018; Katz et al. 2021; Kyle et al. 2022; Nordstrom et al. 2022; Bell et al. 2024). Our objective is to advance such methods with the use of Kompetitive Allele-Specific PCR (KASP) to map putative invasive hybridization. For this, we aim to optimize species-specific markers for the use of eDNA sampling within terrestrial environments, by first testing these markers on blood samples from native, invasive and hybrid individuals, and subsequently testing these approaches both ex-situ (zoo enclosure) and in-situ (St. Eustatius). Additionally, we investigate the dynamics of terrestrial eDNA, including different sampling approaches and substrates, aiming to enhance our understanding of its suitability as a detection tool.

## Methods

To address our research objectives, we developed species-specific KASP assays. KASP involves florescent allele-specific amplification through which species-specific Single Nucleotide Polymorphisms (SNP) variants (homozygous: parental species; heterozygous: hybrids) are determined with two allele-specific primers, requiring no sequencing (Semagn et al. 2014). Testing relied on blood sample assays, and we conducted eDNA sampling both ex-situ at the Rotterdam Zoo and in-situ on the island of Sint Eustatius. Employing surface swabbing and tape-lifting techniques, we collected eDNA samples to assess the efficacy of our assays and compared the levels of positive and negative detection outcomes to determine an optimal sampling strategy.

### Sample collections

To accurately identify and differentiate between pure *I. delicatissima* and hybrids with *I. iguana*, we obtained blood samples derived from previous studies and Rotterdam Zoo collections, all from various genetically confirmed individuals (van den Burg et al. 2018b, 2023) (Appendix I). We included two *I. iguana* specimens from Bonaire and Saba (samples: TBB65, SAB46), three *I. delicatissima* specimens from St. Eustatius (samples: EUX4b, Z18114, Z0208), and one confirmed *I. iguana* x *I. delicatissima* hybrid specimen from St. Eustatius (sample: H3). These samples had been stored at −20°C or −80°C prior to extraction.

To test the efficacy of terrestrial eDNA collections for iguana detections, we first collected different ex-situ samples consisting of sterile swabs (QIAGEN, Germany) and tape-lifts (sterile PCR film, Sarstedt), from an *I. delicatissima* exhibit at the Rotterdam Zoo (Rotterdam, the Netherlands). The exhibit housed two adult *I. delicatissima* individuals, one male and one female. The exhibit itself consisted of an enclosed terrarium located within the Aquarium building of the zoo, kept at a stable 27°C with a relative humidity ranging between 65% and 80%. A misting system was used to maintain the desired humidity, with additional showers (two times a week) with a garden hose. A natural lighting cycle was provided via a transparent ceiling, supplemented with additional UV lights from 7:30 am till 6:30 pm. The interior of the exhibit contained artificial plants and several branches, as well as rock claddings to provide a safe retreat from the viewing window (Fig. 1).

**Fig. 1.**
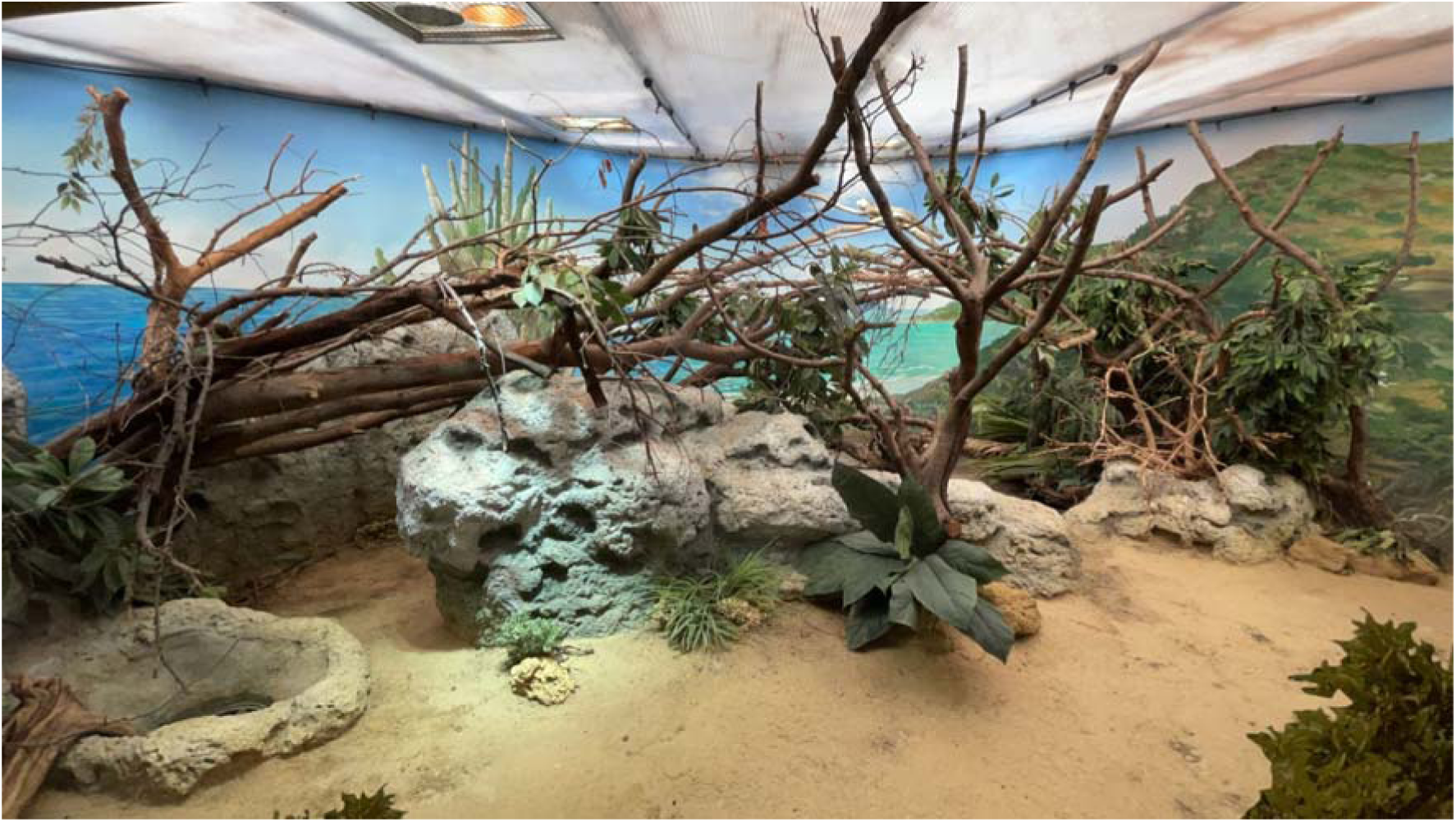
*Iguana delicatissima* exhibit at the Rotterdam Zoo (ex-situ sampling site). Image taken from main viewing window. Image taken by J. van Kuijk.

All exhibit swabs (n=60) were taken from the available environmental surfaces following the sampling protocol of Valentin (2020). A total of 30 dry swabs and 30 wet swabs were taken from three different surfaces: branches, artificial plants, and artificial rock claddings. The swabs were taken with an even pressure by rubbing the swab flat against the aforementioned surfaces in triplicate. The replicates were taken right next to the original sample to minimise the risk of lowering eDNA concentrations by sampling on the exact same spot. A parallel number of 10 wet samples were taken by spraying deionized water on the environmental surface prior to swabbing. Once swabbed, samples were stored in 1 ml Longmire’s Solution (100 mM Tris, 100 mM EDTA, 10 mM NaCl, 0.5% SDS, 0.2% sodium azide) in 2 ml Eppendorf tubes to stabilize the available eDNA, and subsequently stored at −20°C (Longmire et al. 1987). Blank field swabs (both dry and wet) were taken during sampling to track potential contamination.

Sterile PCR plate seals were used as tape-lifts to sample potential eDNA from branches, artificial plants, and artificial rocks in triplicate. The replicates were taken as explained previously. Wet samples (N=5) and dry samples (N=5) were taken of three different surfaces (broad leaves, branches, and rock claddings), where the area for the wet samples was sprayed with deionized water prior to taking the sample. Samples were obtained by wrapping a sterile PCR plate seal tightly over the desired surface to maximize the contact area. Tape lifts were then gently peeled loose and placed in a sterile 50 ml collection tube containing 30 ml Longmire’s solution with sterile forceps.

Opportunistic environmental samples were taken in-situ at two locations on St. Eustatius (Fig. 2). There, upon encountering an iguana (identified as *I. delicatissima* based on morphology) perching on a branch (n=1) or resting on concrete ground (n=1), surface-samples were taken following the protocol of Valentin (2020). Wet and dry swabs were taken in duplicate for both locations, together with a wet and dry negative control during sampling (n=8). Once swabbed, samples were stored in 1 ml Longmire’s in 2 ml Eppendorf tubes to stabilize the available eDNA and placed in a refrigerator on St. Eustatius then stored at −20°C in the Netherlands.

**Fig. 2.**
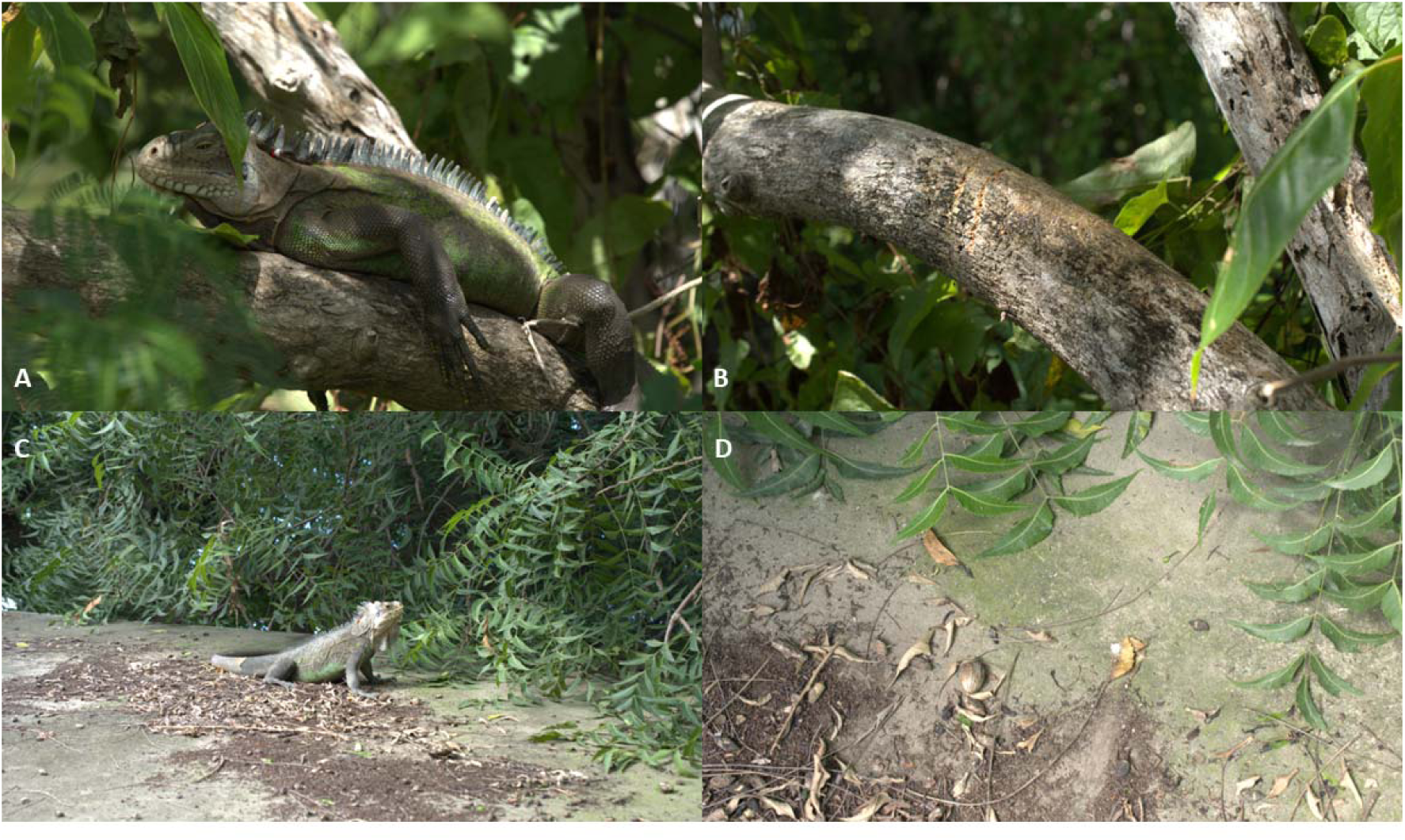
Overview of the in-situ sampled surfaces after a successful iguana encounter. (A) Male *Iguana delicatissima*. (B) Sampled surface consisted of the perching spot (branch) of the iguana in (A). (C) Female *I*. *delicatissima*. (D) Second sampled surface consisted of the resting spot (concrete) of the iguana in (C). Images taken by M.P. van den Burg.

### Sample extraction

DNA from all samples, blood as well as ex-situ and in-situ eDNA samples, were extracted via the isopropanol DNA extraction protocol from the Promega Wizard® Genomic DNA purification kit. Sample material was digested in 300 μl of Nuclei Lysis Solution (Promega) and 3.0 μl Proteinase K. Proteins were precipitated with 100 μl Protein Precipitation Solution (Promega). The supernatant was transferred to a new microcentrifuge tube and 300 μl of 100% isopropanol (kept at −20°C) was added to precipitate the DNA pellet. The DNA pellet was washed with 300 μl 70% ethanol and resuspended in 100 μl TE buffer (Fischer). To give us an estimate of how much general eDNA is collected during sampling, total eDNA concentrations were estimated using a Nanodrop (ND-1000 UV-Vis Thermo Scientific) and aliquoted as undiluted and diluted to 1:10, 1:20, 1:50, and 1:100 concentrations for testing in the KASP assay (see Primer Optimization).

### Primer development

We used published and unpublished genetic data from across the Iguana range to assess the presence of species-diagnostic SNPs for several markers and designed small (<150bp) nuclear eDNA primers for downstream analysis (Stephen et al. 2013; Malone et al. 2017; van den Burg et al. in prep.). We focused on a subset of four nuclear markers: polymerase alpha catalytic subunit (PAC), neurotrophin subunit 3 (NT3), MutL homolog (MLH3), and the oocyte maturation factor (C-MOS) (Appendix II). Sequences were taken from GenBank (Accession numbers in Appendix II). Species diagnostic SNPs were determined by aligning and checking the sequences in Geneious Prime (version 2022.2).

The program Kraken (LGC, Biosearch Technologies) was then used to design two forward primers carrying the standard FAM and HEX tails, with the targeted SNP at the 3′ end, and a common reverse primer. The HEX tail is connected to the *I. delicatissima* SNP, whereas the FAM tail is connected to the *I. iguana* SNP. The SNP sequences from the desired markers with 15 bp left-flanking sequence and 14 bp right-flanking sequence of each SNP site were used in this design.

### Primer optimization and sample testing

A total of four KASP markers were tested on whole blood samples (Appendix I) in 384 well plates with 3µl KASP primer mix and 3 µl DNA template volume. PCR reactions were performed in a HydroCyclyer (LGC, Biosearch Technologies) using the following program: initial denature at 94°C for 15 min, 10 touchdown cycles of 94°C for 20 sec and 61–55°C (dropping 0.6°C per cycle) for 1 min; followed by 26 cycles of 55°C. Plates were read using the PHERAstar plate reader (BMG labtech), and recycled in the thermocycler with 3 additional annealing cycles of 57°C.

All ex-situ and in-situ samples were tested and optimized in 384 well plates with a similar reaction set up of 3 µl KASP primer mix and different DNA template volumes (3 µl, 6 µl, and 9 µl). As KASP amplification has never before been performed on eDNA samples, we used this opportunity to optimize template volume for an effective assay (i.e. to amplify enough nuclear DNA for our SNPs). We were also unsure of the impact of potential inhibitory substances, and thus further tested our samples at five different dilutions; pure, 1:10, 1:20, 1:50, 1:100. PCR reactions were performed using the following program: initial denature at 94°C for 15 min, ten touchdown cycles of 94°C for 20 sec and 61–55°C (dropping 0.6°C per cycle) for 1 min; followed by 26 cycles of 55°C. Plates were read using the PHERAstar plate reader (BMG labtech), and recycled in the thermocycler with four additional annealing cycles of 57°C.

### Data analysis

Data was imported from the PHERAstar plate reader and visualized in RStudio (RStudio team, 2023) using ggplot2 (Wickham 2016) and Cowplot (Wilke 2020). The statistical analysis to determine differences between the sampling methods was done via *t*-tests due to this dataset demonstrating normality via Shapiro-Wilk’s test. Comparisons between sample surfaces were made by testing for normality using Shapiro-Wilk’s test. Further downstream analysis was achieved via the Kruskal-Wallis test, selected due to non-normal data distribution, with a post hoc Dunn’s test from the FSA package (Ogle 2018) to determine differences between sample surfaces. The obtained *p*-values from the Kruskal-Wallis tests were adjusted to account for multiple testing via the benjamini-hochberg procedure.

## Results

### Collected DNA

All together, we were able to collect eDNA from both methods, both ex-situ from the zoo exhibit and in-situ from St. Eustatius. The average total eDNA concentration per sample method showed that ex-situ swabs yielded an eDNA concentration of 15.33 ng/µl ± 19.55 and the ex-situ tape-lift samples yielded 20.2 ng/µl ± 20.71, whereas the in-situ swabs yielded 21.67 ng/µl ± 20.37 (Fig. 3). It should be noted that Nanodrop concentration quantification does not discern whether it is measuring iguanid DNA or DNA from other organic sources. A one-way ANOVA did not reveal a significant difference between the total eDNA concentration and the different methods (F_2, 75_ = 0.552, *p* > 0.05).

**Fig. 3.**
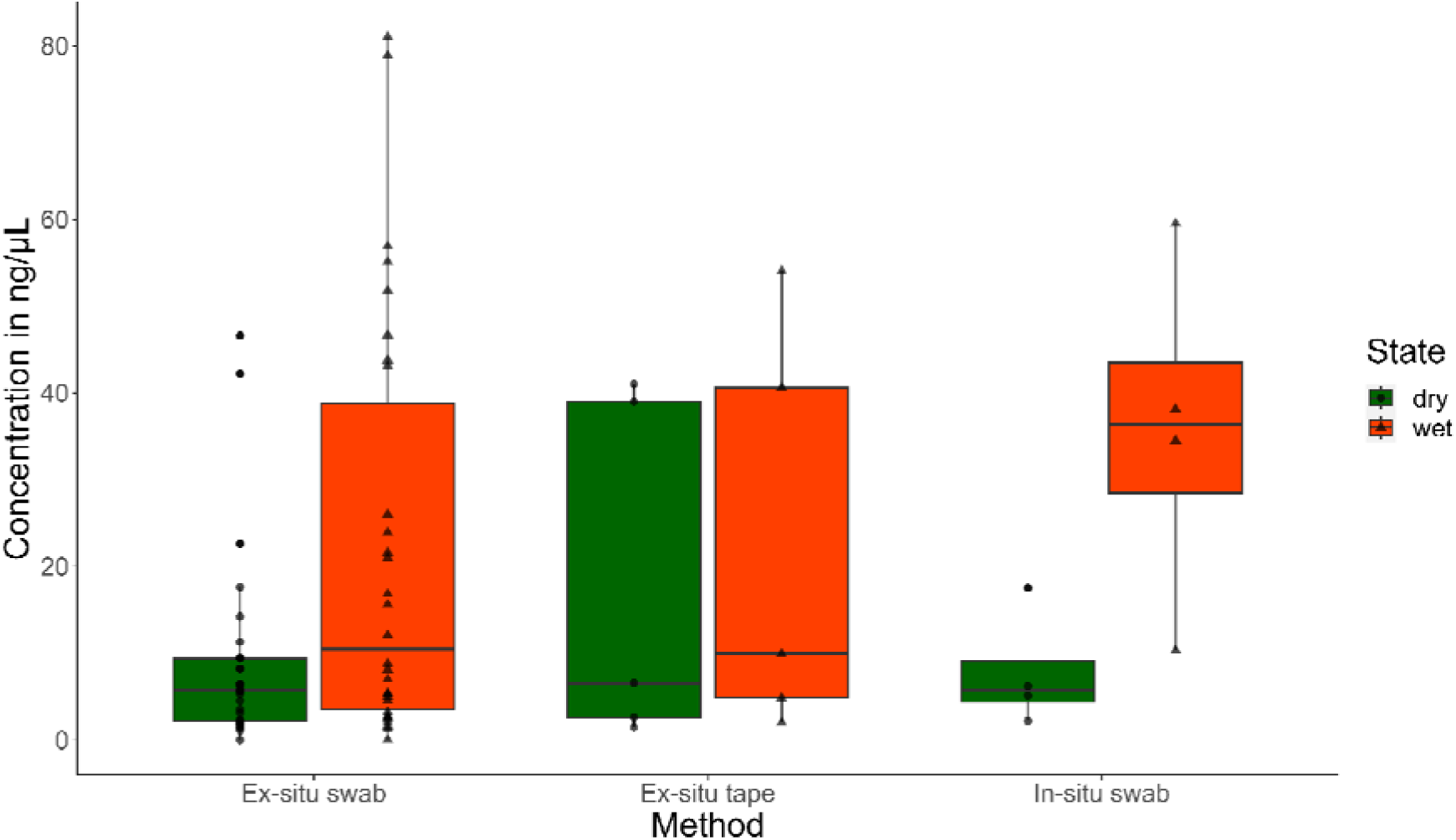
Overview of the average total DNA concentration per sampling method in ng/µl, measured via Nanodrop.

### KASP interpretation

For both ex-situ and in-situ tests, we used positive control samples from known *I. delicatissima*, *I. iguana* and hybrid individuals to ensure accuracy and reliability of our results. These control samples were accepted if they produced a signal that reached over a value of 1.0 on the KASP fluorescence graph, which was indicative of a strong, reliable match, with clusters based on allele-specific fluorescence signals. A result was considered positive for *I. delicatissima* when the fluorescence signal closely clustered to that of the control samples. This method allowed us to confidently differentiate between *I. delicatissima* and *I. iguana*, minimizing the risk of misclassification. Of the four nuclear genes tested, 10 SNPs in PAC, 17 SNPs in MLH3, 4 SNPs in NT3, and 2 SNPs in C-MOS, three SNPs were found to be functional in the KASP assays (1 SNP in CMOS-1, 1 SNP in CMOS-2, and 1 SNP in NT3).

### Ex-situ swabs

Ex-situ swabs from the Rotterdam Zoo exhibit were tested with KASP at a sample dilution of 1:20 with 2 different DNA template volumes (6 µl, and 9 µl), which resulted in 13 positive detections of *I. delicatissima* out of 62 swabs (10.5%). No significant correlation between sample total eDNA concentration and positive iguanid detections was found (*R^2^* = 0.01, *F* = 3.142, *df* = 178, *p* > 0.05).

None of the samples tested positive on all three SNPs. The 6 µl template volume showed amplification at both CMOS-1 and NT3 primers (Fig. 4). The CMOS-2 assay was rejected from further analyses on ex-situ swabs due to non-functional positive controls and potential contamination on the 6 µl template volume plate. The 9 µl DNA template volume did not produce any reliable amplifications and was therefore likewise rejected in further analysis (Fig. 4). A *t*-test on the remaining individual amplifications of the 6 µl template DNA run did not reveal a significant difference in the detection of iguanid DNA between wet swabs and dry swabs (*t*-value = 0.331, *df* = 166, *p* > 0.05).

**Fig. 4.**
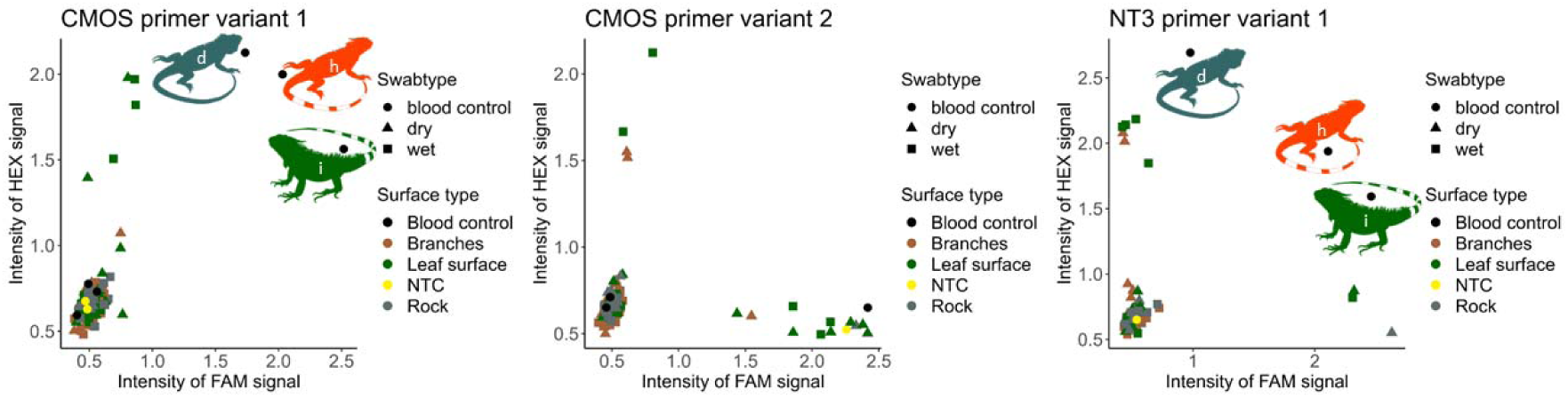
Comparison of the different ex-situ surfaces of the 6 µl template volume, on three different markers from the C-MOS gene and NT3 gene. Surfaces and Swab type did not have a significant influence on the strength of the HEX signal. (d) *Iguana delicatissima*, (i) *I*guana *iguana*, and (h) *delicatissima* x *iguana* hybrid. The NTC stands for the non template control to detect potential contaminations.

A Kruskal-Wallis multiple comparison was conducted to compare the effects of different surfaces on the HEX signal strength. The data included samples tested on three surfaces (Branches, Leaf surfaces, Rocks) and two markers (CMOS-1 and NT3) (Fig. 4). A threshold was set at a HEX signal strength of 1.0, removing all samples below 1.0 that did not show amplification. No significant difference between the three sampled surfaces was found (*X*^2^ = 4.1511, *df* = 2, *p* > 0.05).

### Ex-situ tape-lift samples

Three different surfaces (Branches, Leaf surfaces, and Rocks) were tested for iguana eDNA via tape-lifting, with ten PCR seals as sampling medium, and compared to determine the best sampling surface (Fig. 5). A similar threshold was set at a HEX signal strength of 1.0, removing all samples below 1.0 that did not show amplification. The 6 µl DNA template volume resulted in at total of 14 positive detections (28%), with two samples that tested positive on all three SNPs (samples: TD1, TD2) while tested with undiluted samples. It is important to note that the ex-situ tape-lift CMOS-2 assay included functional positive controls and was therefore incorporated. No significant correlation between sample total eDNA concentration and positive iguanid detections was found (*R^2^* = 0.03, *F* = 2.749, *df* = 147, *p* > 0.05).

**Fig. 5.**
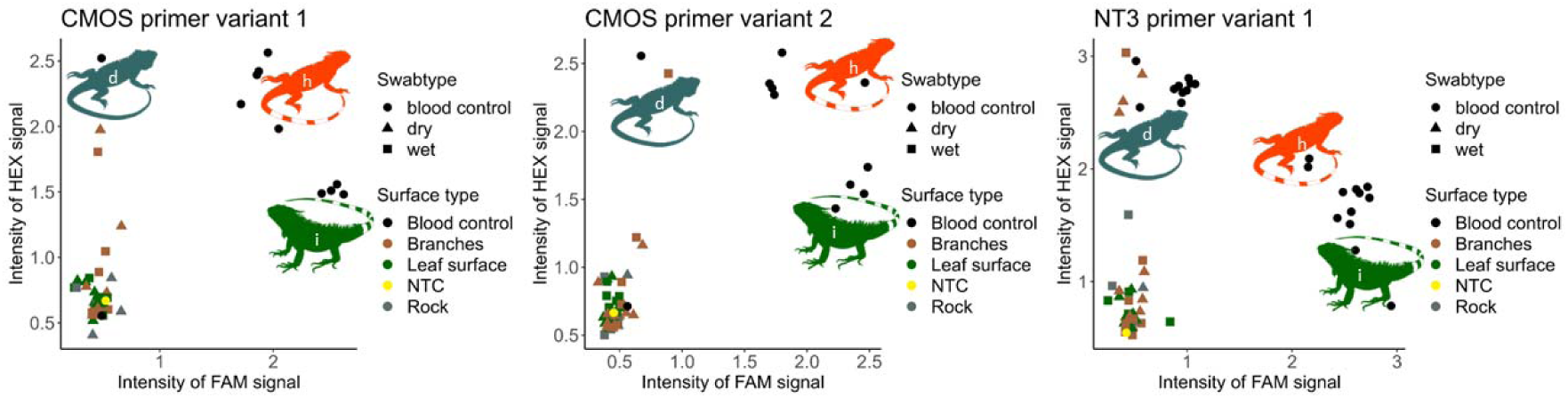
Comparison of the different ex-situ surfaces sampled, combined with a comparison of sample types (wet/dry tape lifts), tested on three different markers from the C-MOS gene and NT3 gene. (d) *Iguana delicatissima*, (i) *I*guana *iguana*, and (h) *delicatissima* x *iguana* hybrid. The NTC stands for the non template control to detect potential contaminations.

A Kruskal-Wallis multiple comparison was used to compare the three sampled surfaces (*X*^2^ = 18.087, *df* = 2, *p* < 0.001). A post hoc Dunn’s Test revealed a significant increase of on branches compared to the other surfaces after multiple testing correction (benjamini-hochberg, *p* < 0.05). A *t*-test revealed no significant difference in the detection of iguanid eDNA between wet swab and dry swab sampling (*t*-value = 0.852, *p*-value > 0.05, *df* = 31).

### In-situ samples

A total of eight different in-situ swabs were sampled on the island of St. Eustatius. The 9 µl DNA template volume showed high amplification with the pure samples across all SNPs (Fig. 6) and resulted in a total of 31 positive detections (77.5%), with 6 undiluted samples tested positive across all three SNPs (samples: 1Wa, 1Wb, 1Da, 1Db, 2Da and 2Db). It is important to note that the in-situ CMOS-2 assay included functional positive controls and was therefore incorporated. A significant correlation between sample total eDNA concentration and positive detections was found (*R^2^* = 0.18, *F* = 13.02, *p* < 0.05, *df* = 147). The outcomes of the two sampled surfaces (Branches, Concrete) were compared with a Kruskal-Wallis multiple comparison to determine a difference in outcomes. A threshold was set at a HEX signal strength of 1.0 to remove all samples below that did not show amplification. No significant difference between the sampled surfaces was found (*X*^2^_8,1_ = 0.0369, *p* > 0.05). Swab types were also compared with a *t*-test to determine differences in swab types. No significant difference in the detection of iguanid eDNA between wet and dry swab sample collection was found (*t* = −1.688, *df* = 147, *p* = > 0.05).

**Fig. 6.**
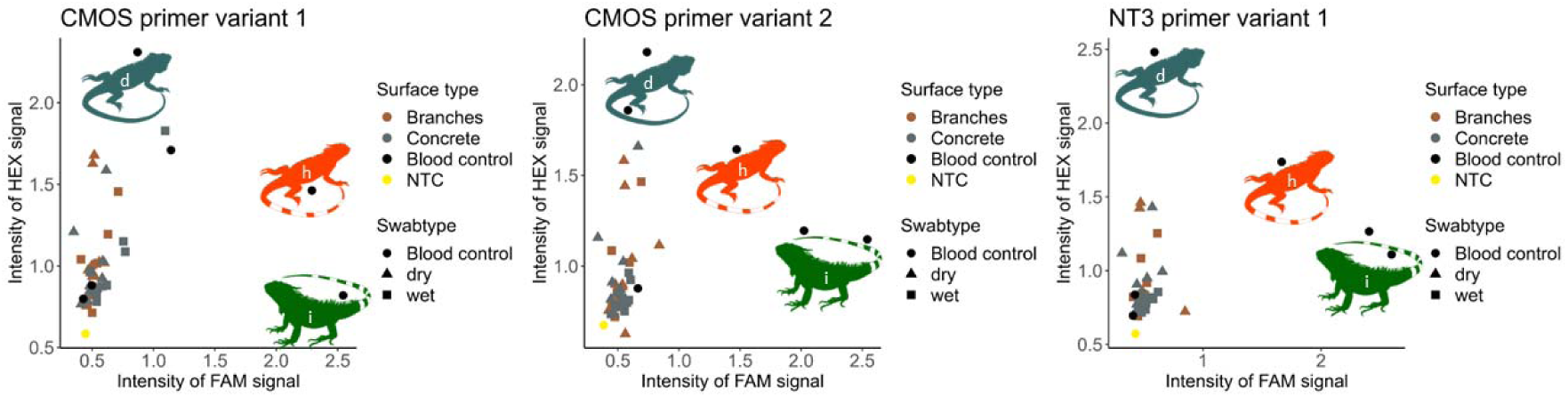
Comparison of the different in-situ different surfaces, combined with a comparison of sample types (wet/dry swabs), tested on three different markers from the C-MOS gene and NT3 gene. (d) *Iguana delicatissima*, (i) *Iguana iguana*, and (h) *delicatissima* x *iguana* hybrid. Swabs were taken from the surface on which the iguana was encountered. The NTC stands for the non template control to detect potential contaminations.

### Overview

In summary (see Table 1), we found that 16,1% of the ex-situ swabs resulted in positive detections for two SNPs (the third SNP marker failed), tested with a templated volume of 6 µl and a dilution of 1:20. Within the ex-situ tape-lift samples, we found that 28% resulted in positive detections, when tested with a 6 µl template volume and in a 1:20 dilution. Finally, we found that 77.5% of the in-situ samples resulted in positive detections for all three SNPs, while testing undiluted samples with a template volume of 9 µl. Furthermore, a significant correlation was found between the total eDNA concentration of the in-situ samples and the number of positive detections.

**Table 1.** Summary of the results from the different sample techniques used in this study. Included are the optimal DNA template volume, optimal sample dilution, and sample success rate for each sampling technique.

| Summary table | In-situ swabs | Ex-situ swabs | Ex-situ tape-lift |
| --- | --- | --- | --- |
| **Optimal template volume** | 9 μl DNA template | 6 μl DNA template | 6 μl DNA template |
| **Optimal dilution** | Undiluted samples | 1 :20 | 1:20 |
| **Markers** | Cmos1, Cmos2 & NT3 | Cmos1, NT3 | Cmos1, Cmos2 & NT3 |
| **Percentage positive detections** | 77.5% | 16.1 % | 28.0% |

## Discussion

Here, we develop a new, non-invasive approach for the detection of invasive species and putative invasive hybridization in iguanas using terrestrial eDNA. Specifically, we designed species-specific, nuclear SNPs, and examined different eDNA sampling strategies as well as investigated different substrate locations for terrestrial eDNA. Our results show that we are able to successfully detect the presence of *I. delicatissima* in ex-situ and in-situ samples collected from both swabs, and tape-lifting. This study thus provides the first proof-of-concept evidence that eDNA, a novel cost-effective and efficient sampling method, can be utilized to survey terrestrial habitats across different substrates for iguana presence, as well as possible invasive hybridization. Notably, this is also the first study (to our knowledge) to combine fast and low-cost KASP analysis to eDNA collections, helping to process and expedite bi-allelic nuclear data, a necessary first-step in detecting potential hybrid individuals.

The ability to detect rare non-native animals and their hybrids, or rare surviving native animals, is especially critical at different spectra of the invasion process and for opposite conservation reasons. For example, eradication of invasive animals at the onset of invasion, or the identification and rescue of last native survivors before they too are usurped when invasion proceeds to full extinction of the native populations, is vital for safeguarding biodiversity (e.g., Milinkovitch et al. 2013; Miller et al 2017; Debrot et al. 2022; Pounder et al. 2020). The application of eDNA is thus likely to be of great value for early invasion confirmation, particularly for presumed presence or absence of the target species (Rout et al. 2014; Russell et al. 2017), be it invasive species or rare, threatened native taxa (Stewart et al. 2017). Additionally, eDNA enables the simultaneous monitoring of multiple species from a single sample, offering a comprehensive view of ecosystem health. Of note, using eDNA methods also eliminates the need for CITES permits, significantly speeding-up workflows.

While other terrestrial eDNA studies have used universal mitochondrial markers that have a high success rate due to high copy number compared to nuclear DNA (Valentin et al. 2020; Andres et al 2021; Lyet et al. 2021), we used custom-made nDNA SNP markers derived from previous studies (Malone et al. 2017; van den Burg et al. 2018b). This offered us more detailed insights into an individual’s genotype, allowing for more in-depth genetic analysis and conservation strategies. However, during our experiments, the markers PAC and MLH3 did not function as hoped. This issue may be attributed to the high density of SNPs closely clustered around the SNP of interest, leading to excessive individual variation and primer mismatches (Wilcox et al. 2013). Additionally, the presence of highly similar homoeologous or paralogous sequences could have contributed to these mismatches (Makhoul et al. 2020). As a result, both markers were excluded from the experiments. Furthermore, the C-MOS 2 marker showed inconsistent performance; it did not function in the ex-situ analysis in which we would expect high residual eDNA of interest, but worked as intended in the in-situ analysis. This discrepancy might be due to technical errors or contamination during preparation and processing of the ex-situ plate despite repeated tests. Though this seems unlikely, it certainly warrants further investigation.

These challenges might contribute to the observed low iguana eDNA detection rates in our study. A minority (c. 16%) of swabs taken at the iguana exhibit at the Rotterdam Zoo resulted in positive detection. One explanation for this low eDNA detection rate could be the insufficient presence of nuclear eDNA available on the sampled surface (Adams et al. 2019; Sigsgaard et al. 2020) due to any number of possible biotic and abiotic forces dictating eDNA shedding, transport or degradation (Stewart 2019). An alternative explanation could be the shortage of iguana eDNA on artificial surfaces, or simply the concentration of eDNA in the samples we collected due to methodological shortcomings. Contrary to expectations, this study did not find a significant difference between swab types (wet or dry) and sample surfaces, whereas Valentin (2018) argued that water is the dominant mechanism of eDNA transport even for terrestrial systems. Sample collection via tape-lifting yielded a higher proportion of positive detections (28%) and as far as we know, has not been tested previously. Our findings indicate that branches, which serve as natural perching spots for iguanas, resulted in a significantly higher number of positive detections compared to rocks and leaves, suggesting that species-specific behaviour is important to consider for eDNA sampling. The in-situ samples from St. Eustatius directly taken after spotting an iguana had a considerably higher success rate for positive iguana eDNA detection (77.5%). It should be noted, however, that sampling directly after encounters ensured minimal environmental degradation of the sample DNA.

In this study we found three successful nDNA SNPs that can be applied to genotype and distinguish putative hybrid iguanas via DNA samples derived from blood and eDNA. When comparing sampling methods, we observed no significant difference in swab types used for *Iguana* eDNA detection. In general, we found a tendency for perching spots such as broad branches to exhibit a higher rate of amplification when compared to other surfaces. A possible explanation for this phenomenon is that iguanid lizards mark their territory via waxy lipids released from their femoral pores (Alberts 1990). The femoral pores are holocrine secretory glands located on the thighs and are found in all members of the Iguanidae (Mayerl et al. 2015). Since iguana’s mark perching spots as part of their territory, it is reasonable to assume that DNA is also deposited on these sites. Therefore, sampling perching spots or similar objects in the habitat that are likely to be marked with waxy lipids by iguanas could therefore increase the overall detection of iguanid lizards. Additionally, future studies could assess variation between sex and eDNA deposition as Iguana sexes are known to differ in femoral pore sizes (Bakhuis. 1982).

We demonstrate the viability of detecting *Iguana* eDNA in terrestrial environments, from surfaces and vegetation using various sampling techniques. It should be noted that our markers and approach worked “in silico”, as well as with blood samples from both species and hybrids. However, we were only able to test environmental samples from *I. delicatissima* due to logistical constraints. We would recommend an additional study to test the approach on environmental samples from *I. iguana* and their hybrids. Sampling strategies should focus on the specific microhabitat use and life cycle of the target species (Adams et al. 2019a), and monitoring threatened or endangered species should prioritize the minimizing of potential sampling hazards to the focal organism or habitat. For example, we currently are unable to ascertain how long this *Iguana* eDNA lasts on environmental surfaces. If prolonged detection is possible, we foresee these terrestrial eDNA approaches to be useful in sampling environmental surfaces to prospect for potential invasive or hybrid presence (i.e. sight unseen). This eDNA method could also be valuable in establishing the probability of hybrid individuals that do not demonstrate obvious morphological signs of genetic admixture. This would enable more targeted response measures after eDNA analysis. Additionally, rigorous validation and optimization steps of assays are required to enhance our understanding of any confounding factors that might influence the performance and efficiency of eDNA detection in cryptic taxa.

## Conclusion

This research is relevant to a broad spectrum of terrestrial biodiversity managers and researchers as it provides a practical solution for conservation practitioners, particularly those working to protect the Lesser Antillean Iguana and potentially other cryptic terrestrial species facing similar threats. The implications of our findings extend beyond the Caribbean, offering a methodology that could be adapted for use in other regions and for other species at risk of hybridization or extinction due to invasive species. We believe that our study will be of significant interest given the global relevance of biodiversity conservation and the increasing threat of invasive species. The novel application of terrestrial eDNA sampling techniques for species conservation represents a significant advancement in the field, offering a new tool that can enhance the speed and accuracy of conservation interventions.

## Acknowledgements

Rotterdam Zoo, STENAPA and RAVON provided support and resources during the project. Ton Weber, Sandra Bijhold, Linda Bruins-van Sonsbeek, Kornelia Serwatowska, Bobbie Sewalt, Carola Feijt, Dennis Klein Gunnewiek and Fenna Splinter provided invaluable assistance, feedback, and ideas during the fieldwork and lab analysis. This project was funded by the International Iguana Foundation (IIF) and Wageningen Marine Research (Project BO Soortenbescherming Caribisch Nederland; BO-43-117-006).

## Author Contributions

Authors JLK and KAS contributed to the study conception and design. Material preparation, data collection and analysis were performed by JLK, MPB, EAD, MB and BW. The first draft of the manuscript was written by JLK and all authors commented on previous versions of the manuscript, contributing to writing. All authors read and approved the final manuscript.

## Notes

### Competing Interest Statement

The authors have declared no competing interest.

